# Novel centriolar defects underlie a primary ciliary dyskinesia phenotype in an adenylate kinase 7 deficient ciliated epithelium

**DOI:** 10.1101/2023.07.25.550535

**Authors:** Jennifer Sheridan, Aline Grata, Eve E. Suva, Enzo Bresteau, Linus R. Mitchell, Osama Hassan, Brian Mitchell

**Affiliations:** Northwestern University, Feinberg School of Medicine, Department of Cell and Developmental Biology; Northwestern University, Lurie Cancer Center

**Author notes:** Corresponding author Address: Brian Mitchell, Northwestern University, Feinberg School of Medicine, 303 E. Chicago Ave, Chicago IL 60611. These authors contributed equally (Jennifer Sheridan, Aline Grata).

## Abstract

The skin of *Xenopus* embryos contains numerous multiciliated cells (MCCs), which collectively generate a directed fluid flow across the epithelial surface essential for distributing the overlaying mucous. MCCs develop into highly specialized cells to generate this flow, containing approximately 150 evenly spaced centrioles that give rise to motile cilia. MCC-driven fluid flow can be impaired when ciliary dysfunction occurs, resulting in primary ciliary dyskinesia (PCD) in humans. Mutations in a large number of genes (∼50) have been found to be causative to PCD. Recently, studies have linked low levels of Adenylate Kinase 7 (AK7) gene expression to patients with PCD; however, the mechanism for this link remains unclear. Additionally, AK7 mutations have been linked to multiple PCD patients. Adenylate kinases modulate ATP production and consumption, with AK7 explicitly associated with motile cilia. Here we reproduce an AK7 PCD-like phenotype in *Xenopus* and describe the cellular consequences that occur with manipulation of AK7 levels. We show that AK7 localizes throughout the cilia in a DPY30 domain-dependent manner, suggesting a ciliary function. Additionally, we find that AK7 overexpression increases centriole number, suggesting a role in regulating centriole biogenesis. We find that in AK7-depleted embryos, cilia number, length, and beat frequency are all reduced, which in turn, significantly decreases the tissue-wide mucociliary flow. Additionally, we find a decrease in centriole number and an increase in sub-apical centrioles, implying that AK7 influences both centriole biogenesis and docking, which we propose underlie its defect in ciliogenesis. We propose that AK7 plays a role in PCD by impacting centriole biogenesis and apical docking, ultimately leading to ciliogenesis defects that impair mucociliary clearance.

## Introduction

Cilia, both nonmotile and motile, are highly conserved microtubule structures that play an essential role in numerous physiological processes (Anvarian et al., 2019, Bustamante-Marin and Ostrowski, 2017, Reiter and Leroux, 2017). Motile cilia can form singularly, as in sperm flagella, or in the cells at the left-right organizer of the embryonic node. Alternatively, numerous cilia can form in a complex cell known as a multiciliated cell (MCC), whose cilia coordinately and rhythmically beat to generate a directed fluid flow (Boutin and Kodjabachian, 2019, Brooks and Wallingford, 2014, Meunier and Azimzadeh, 2016). Proper MCC development and function depends on a multistep process involving centriolar amplification, producing dozens of centrioles or basal bodies, which migrate apically and dock at the apical surface and give rise to motile cilia (Figure 1A). MCCs are found in numerous tissues where their role in creating a directed fluid flow is necessary, such as in the brain ependyma and in the fallopian tubes (Spassky and Meunier, 2017). Furthermore, MCCs line the respiratory epithelium, where their motile cilia beat in a coordinated manner to drive directed mucociliary flow (Bustamante-Marin and Ostrowski, 2017). A loss in the directed fluid flow derived from motile cilia defects leads to a condition known as Primary Ciliary Dyskinesia (PCD) (OMIM 244400) (Bush, 2000, Bustamante-Marin and Ostrowski, 2017, Legendre et al., 2021).

**Figure 1.**
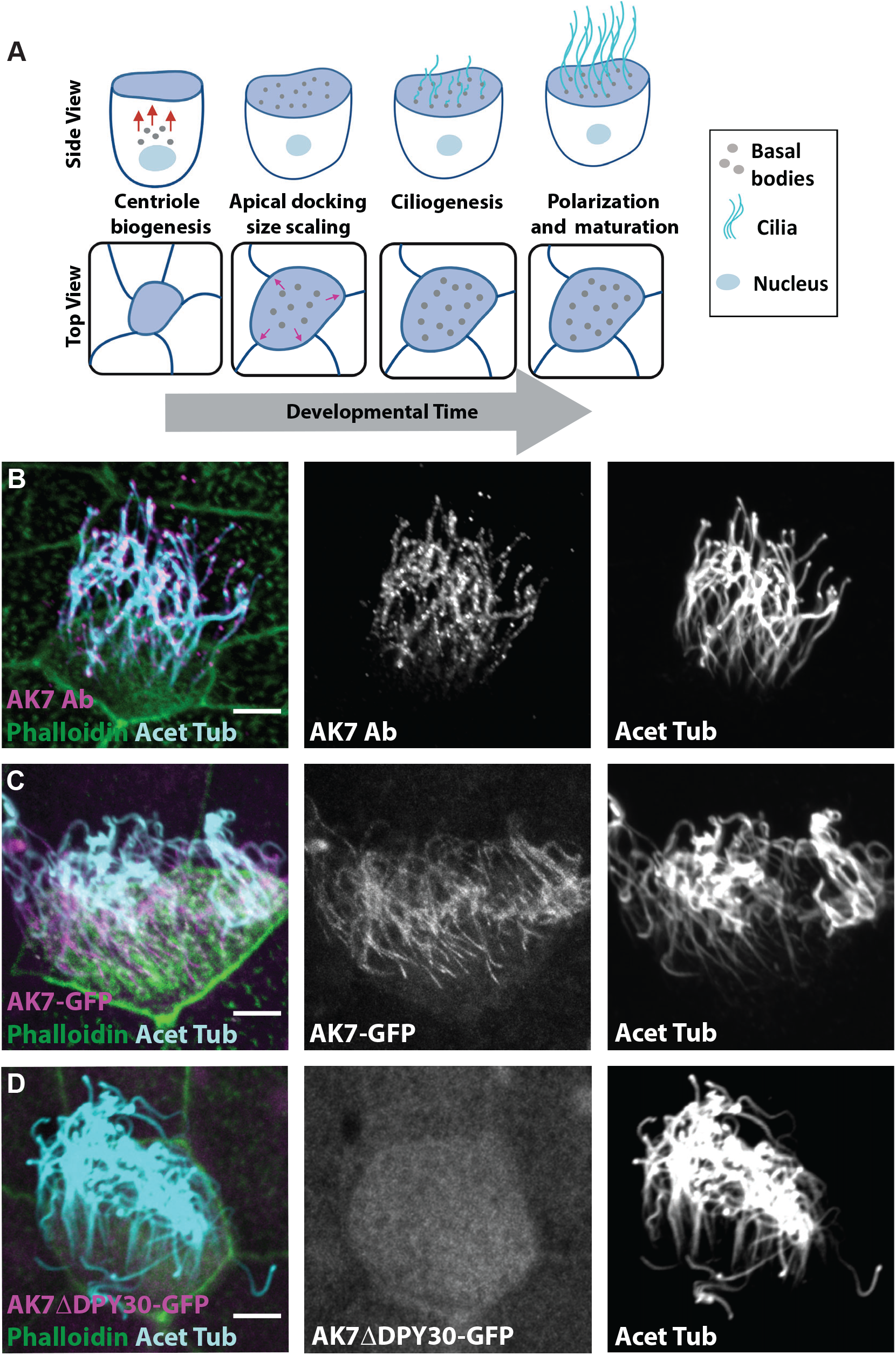
Localization of AK7 to cilia. (A) Schematic representing MCC development illustrating the processes of centriole biogenesis, apical docking and MCC’s apical area expansion and ciliogenesis. (B) Representative image of AK7 ciliary localization in MCC stained with anti-AK7 (magenta and white) and anti-acetylated tubulin (cyan and white) antibodies together with phalloidin (green). (C) Representative image of a MCC from an embryo injected with AK7-GFP (magenta, white) and stained with anti-acetylated tubulin (cyan, white) antibody together with phalloidin (green). (D) Representative image of a MCC from an embryo injected with AK7ΔDPY30-GFP (magenta, white) and stained with anti-acetylated tubulin (cyan, white) antibody together with phalloidin (green). Scale bars are 5 μm.

PCD is a rare inherited condition characterized by impaired ciliary function resulting from genetic mutations (Goutaki and Shoemark, 2022, Legendre et al., 2021, Lucas et al., 2020, Zariwala et al., 2019, Bush, 2000). Due to the wide distribution of motile cilia in diverse tissues, PCD presents with a broad range of symptoms, including respiratory dysfunction, *situs inversus*, infertility, and in rare cases, hydrocephaly. PCD has a complex etiology, in part due to the complex nature of the ciliary molecular machine that includes hundreds or thousands of distinct proteins (McCafferty et al., 2023). PCD was initially classified by genetic mutations producing defects in ciliary ultrastructure that is observed as defects in axonemal dynein arms; however, now approximately 50 genes have been identified as causative in PCD patients (Horani and Ferkol, 2021, Zariwala et al., 2019). Some of those genes are distinct from classical PCD genes, such as *MCIDAS, CCNO, FOXJ1*, and *GAS2L2*, which regulate MCC differentiation, centriole amplification, ciliogenesis, and cilia polarity, respectively, yet are all associated with an impaired directed fluid flow (Amirav et al., 2016, Pan et al., 2007, Stubbs et al., 2008, Stubbs et al., 2012, Yu et al., 2008, Boon et al., 2014, Bustamante-Marin et al., 2019, Horani and Ferkol, 2021, Wallmeier et al., 2014, Zariwala et al., 2019). Notably, up to 30% of individuals with PCD have no identified pathogenic mutations in related genes, suggesting that, genetically, we still have significantly more to learn about PCD (Zariwala et al., 2019).

Adenylate kinases are metabolic enzymes that catalyze the reaction of 2 ADP ↔ 1 AMP + 1 ATP (Dzeja and Terzic, 2009, Otokawa, 1974). They are important modulators of cellular energy levels and have additionally been linked to AMP signaling via AMP Kinase. Mutations in one member of the family of adenylate kinases, adenylate kinase 7 (AK7), have been identified in several patients with PCD (Mata et al., 2012). Furthermore, mice deficient for AK7 produce PCD phenotypes such as ciliary ultrastructure defects and impaired ciliary beat frequency (Fernandez-Gonzalez et al., 2009). Interestingly, low levels of AK7 have been found to correlate with decreased beat frequency within broad cohorts of patients, suggesting that even in patients with a normal AK7 gene, there may be feedback loops that lead to dysregulation of AK7 expression or protein stability (Mata et al., 2012, Milara et al., 2010). Despite these numerous links between AK7 and PCD, there remains a fundamental gap in our understanding of AK7’s role in MCC function. Motile cilia depend heavily on copious energy supplies for proper beating, which could require the function of AK7 (Dzeja and Terzic, 2009, Otokawa, 1974). Alternatively, AK7 could be important for numerous aspects of MCC development. In this study, we examined AK7 CRISPR/Cas9 generated mutations in *Xenopus*, which presented with PCD phenotypes of impaired tissue-wide directed fluid flow and a mild decrease in ciliary beat frequency. Importantly, our findings indicate that AK7 has multiple roles in proper MCC formation and function.

## Results

### AK7 localizes along cilia of MCCs

AK7 is highly associated with ciliated epithelial tissues and has been shown to localize at the apical surface of ciliated cells and to sperm flagella (Lores et al., 2018, Milara et al., 2010). To explore AK7 localization in the *Xenopus* ciliated epithelium, we performed immunofluorescent (IF) staining using an AK7 antibody together with the standard cilia marker, acetylated tubulin. Stage (ST) 28 embryos contain a mature ciliated epithelium and we found co-localization of AK7 with acetylated tubulin in the cilia (Figure 1B). Furthermore, this localization was confirmed using injection of mRNA encoding a fluorescently tagged version of the full length AK7 (AK7-GFP) (Figure 1C). Importantly, unlike other AKs, AK7 has as a DPY30 domain (Lores et al., 2018). DPY30 domains have previously been identified in multiple cilia proteins and are known to be important for the ciliary function of RSP2 (Gopal et al., 2012). Additionally, DPY30 domains share similarity to the RIIa domain known for dimerization and docking of the cyclic AMP-dependent protein kinase and A kinase anchoring proteins (AKAPs) (Kinderman et al., 2006, Gopal et al., 2012). We generated a version of AK7 lacking the DPY30 domain and found that the ciliary localization is dramatically reduced, suggesting that this domain is important for its ciliary localization (Figure 1D). The localization of AK7 to cilia suggests its function could be regulating ATP levels for the energetically taxing motile cilia or potentially to facilitate signaling through cAMP.

### AK7 depleted embryos show an impaired mucociliary fluid flow

AK7 expression levels have been found to correlate with ciliary function in patients with PCD (Milara et al., 2010). However, despite these links, a causative understanding of the ciliary defects produced by depletion of AK7 remains largely unexplored. To investigate and characterize this, we performed CRISPR/Cas9-mediated editing of *AK7*, combining two guide RNAs (gRNAs) targeting exons 1 and 4 of the gene (Supplementary (S) Figure S1A). The efficacy of the gRNAs on the target DNA was assayed *in vitro* (Figure S1B) (Bhattacharya et al., 2015, Wiedenheft et al., 2012). *Xenopus* embryos were injected with the two guide RNAs and Cas9 protein for the knockout condition (AK7-KO or AK7 Crispant), while Cas9 alone or gRNAs alone were used as controls (hereafter referred to as Cas9 and gRNAs, respectively). Importantly, in AK7 Crispant embryos we find a dramatic loss of AK7 antibody staining (Figure 2A). We next investigated possible PCD-related phenotypes in the AK7 Cispant embryos. PCD is mainly characterized by an impaired mucociliary function, hence, we tested the generation of directed flow in control and AK7 Crispant embryos. The *Xenopus* embryonic epithelium is covered in mucus, which moves along the epithelial surface in a flow sustained by the coordinated ciliary beating of MCCs. We applied fluorescent microspheres on the anterior side of anesthetized and immobilized live embryos at ST28 when directed flow is robust (Werner and Mitchell, 2013). In contrast to control embryos which generated vigorous flow, we observed a significant decrease in fluid flow in the AK7 Crispants (Figure 2B). Importantly, fluid flow returned to normal levels when we performed a rescue experiment by co-injecting embryos with AK7 mRNA not targeted by the gRNAs (Figure 2B). This result confirms the importance of AK7 in tissue level mucociliary clearance and is consistent with a role in PCD.

**Figure 2.**
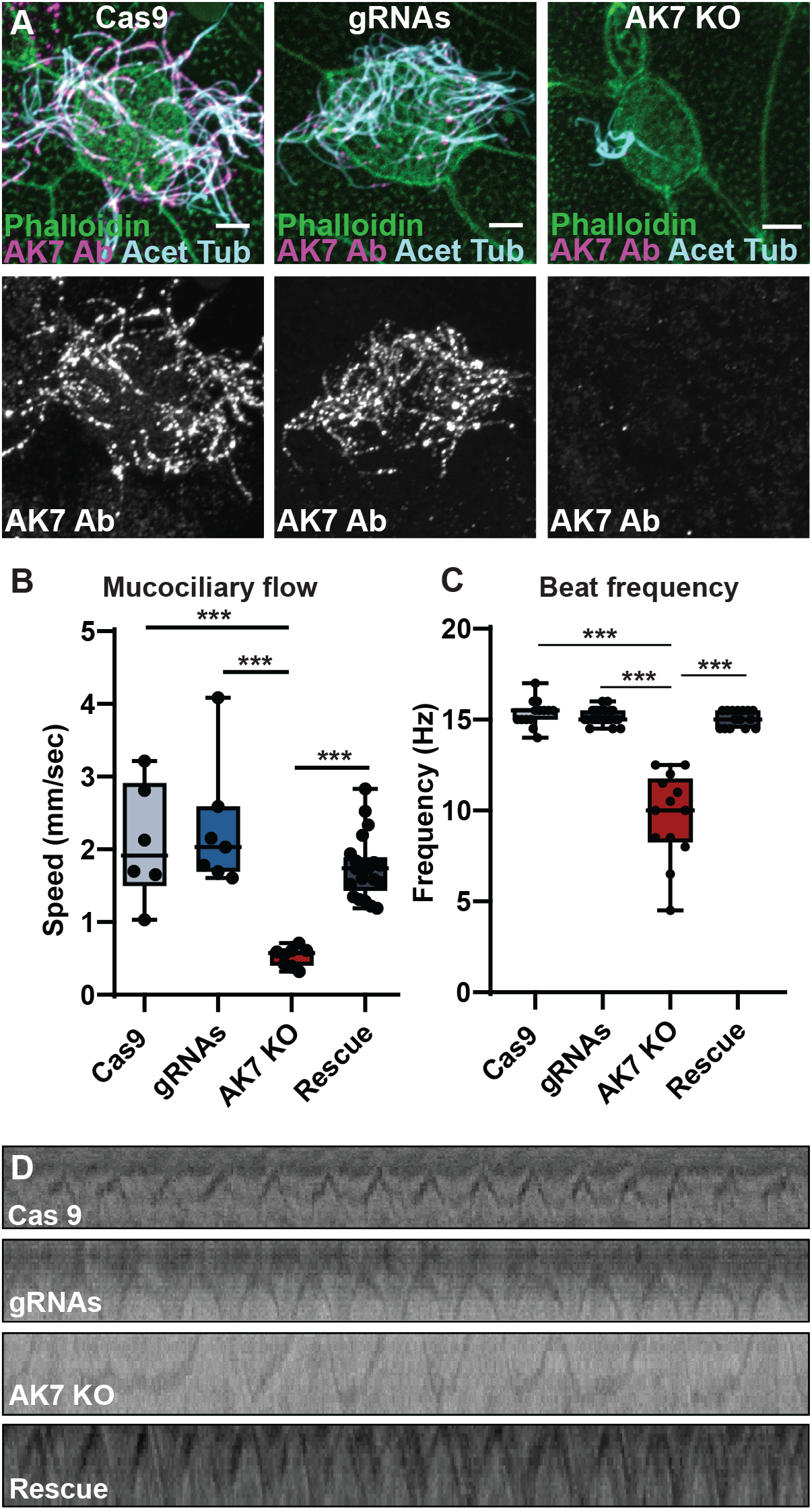
AK7 deletion causes an impairment of fluid flow and ciliary function. (A) Representative images of MCCs stained with anti-AK7 (magenta, white) and anti-acetylated tubulin (cyan) antibodies together with phalloidin (green) showing strong ciliary staining in controls (Cas9 alone or gRNA alone) that is missing in AK7 KO cells (Cas9 + gRNAs). Scale is 5 μm. (B) Quantification of mucociliary flow using the displacement of fluorescent beads in embryos injected with Cas9 alone (n = 3 embryos, 6 flow measurements), gRNA alone (n = 3 embryos 7 flow measurements), Cas9 + gRNA (AK7 KO; n = 2 embryos, 8 flow measurements), or Cas9 + gRNA + AK7-GFP mRNA (Rescue; n = 3 embryos 21 flow measurements). (C) Quantification of ciliary beat frequency in Cas9 alone (n = 3 embryos, 14 cells), gRNA alone (n = 3 embryos, 24 cells), Cas9 + gRNA (AK7 KO; n = 5 embryos, 13 cells), or Cas9 + gRNA + AK7-GFP mRNA (Rescue; n = 4 embryos, 15 cells). (D) Representative kymographs of 1 second from our high-speed movies used to quantify beat frequency (See Supplemental Movies 1-4). Graphs in B and C are box and whisker plots where the error bars represent the min and max values.

### AK7 depleted MCCs have impaired ciliary beating

The loss of tissue level flow can be due to multiple underlying defects, including loss of MCCs, loss of cilia, or loss of ciliary motility. To begin to characterize the cellular level defects in AK7 Crispant embryos, we performed live imaging of ciliary beating. We used high-speed light microscopy to capture MCCs’ ciliary movements (Supplementary Movies 1-4). We quantified beat frequencies in the KO and control conditions using kymographs and found that that there was a clear decrease in beat frequency in the AK7 KO embryos (Figure 2C and 2D). Healthy wild type MCCs have a beat frequency of about 15 Hz (Werner et al., 2013), similar to the rate that we measured in our control groups (15.3 ± 0.7 Hz and 15.7 ± 0.4 Hz in the Cas9 or gRNAs conditions, respectively). In contrast, AK7-KO MCCs had a significant drop in beating frequency, with an average of 9.7 Hz ± 2.4 Hz (Figure 2C and D). These results align with previous papers in other model systems that reported a decreased beat frequency in AK7 knockouts and mutants (Fernandez et al., 2009; Milara et al., 2010). However, the roughly 30% decrease in beat frequency, while significant, did not seem consistent with the almost complete loss of fluid flow that we observed, suggesting that there could be additional phenotypes associated with the loss of AK7.

### AK7-depleted MCCs display defects in cilia number and morphology

Our results indicate that a loss of AK7 expression in the *Xenopus* ciliated epithelium impairs mucociliary clearance and ciliary beating. To address the morphological phenotypes of AK7 depletion on cilia, we performed both confocal and scanning electron microscopy (SEM) imaging. We performed IF on ST28 embryos using anti-acetylated tubulin antibodies to visualize cilia (Figure 3A). In control embryos MCCs display an average of 150-200 cilia. Given the difficulty in precisely counting cilia number in single MCCs, we quantified the gross amount of cilia per cell by dividing cells in three categories: cells with more than 50 cilia (corresponding to what is expected in wildtype conditions), cells with 10 to 50 cilia, and cells with less than 10 cilia. In our control conditions, almost all MCCs had more than 50 cilia (96.3% in gRNAs and 100% in Cas9, Figure 3C). Noticeably, in AK7 Crispant MCCs, we observed no MCCs with more than 50 cilia, while 55% of cells had less than 10 cilia, and the remaining 45% had 10 to 50 cilia (Figures 3A and 3C). These defects in cilia number are also observable in SEM images of ST28 embryos (Figure 3B and Figure S2B). Furthermore, we measured the length of cilia using our side view imaging of cilia beating and found that cilia length decreased from ∼ 15 mm in controls animals to 11mm in AK7 Crispants (Figure 3D and S2A). While significant, this analysis selected for only motile cilia and thus likely represents an underestimation of the overall effect on ciliary length. These results are consistent with the effects observed in SEM images (Figure 3B and Supplementary Figure S2B). Importantly, we can rescue the ciliary length and number defects by overexpressing full length AK7 (Figure 3C-D). These results suggest that in addition to the defects in cilia motility reported above, AK7 deficient cells also display a significant impairment in ciliary biogenesis and morphology. This defect in ciliogenesis is an important component to the overall loss of ciliary clearance and flow.

**Figure 3:**
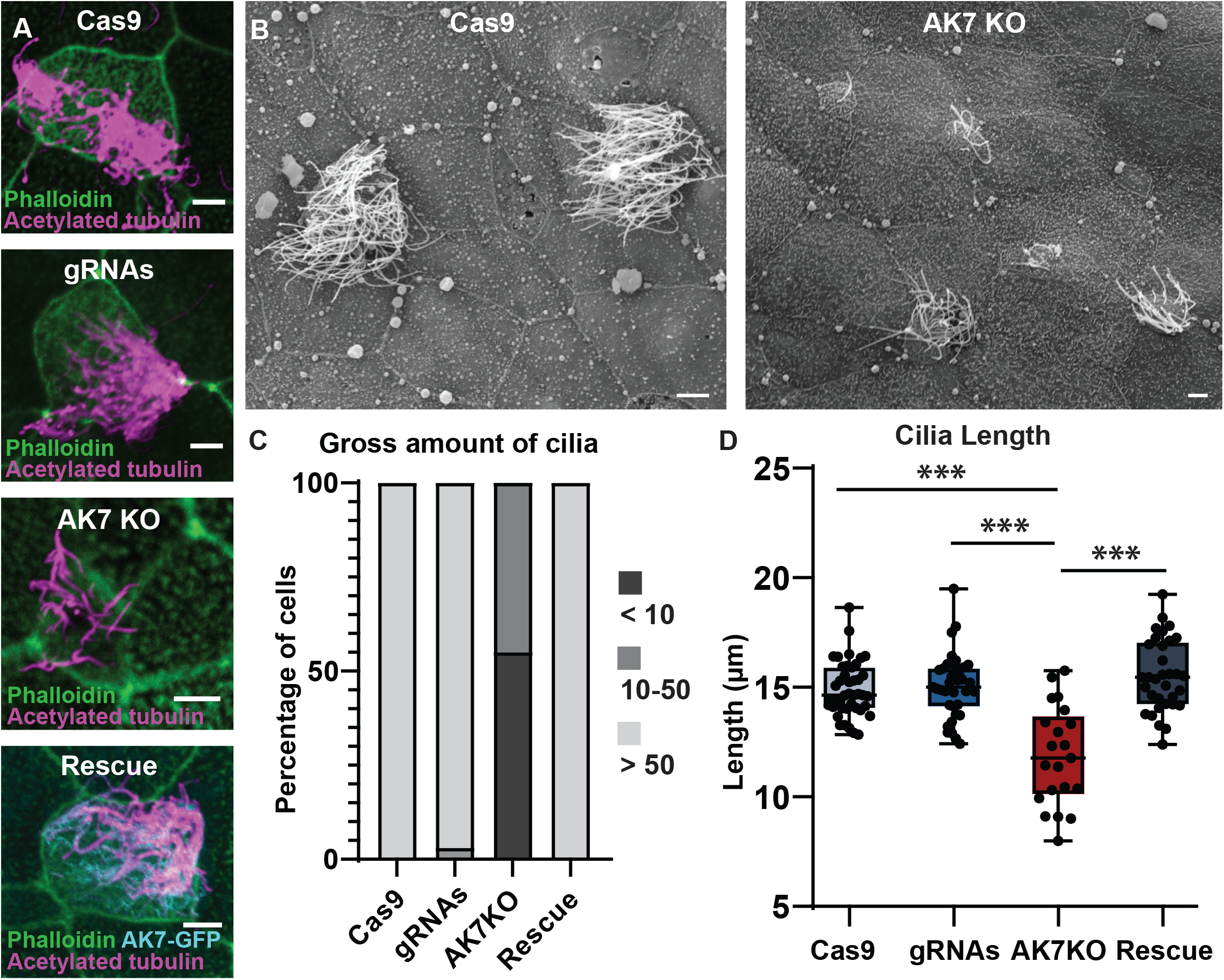
AK7 deletion causes ciliogenesis defects. (A) Representative images of control (Cas9 and gRNA), AK7 KO and Rescue cells stained with anti-acetylated tubulin (magenta), AK7-GFP (cyan) and phalloidin (green). (B) Representative SEM images of control (Cas9 alone) and AK7 KO embryos. Scale in (A) and (B) is 5 μm. (C) Binned quantification of cilia number scoring cells as having > 50, 10-50 or < 10 cilia from embryos injected with Cas9 alone (n = 5 embryos, 26 cells), gRNA alone (n = 7 embryos, 28 cells), AK7 KO (n = 5 embryos, 20 cells), and Rescue (n = 7 embryos, 26 cells). (D) Quantification of cilia length taken from side projection movies of cilia beating (n = 6 embryos, 42 cilia (Cas9), 6 embryos, 35 cilia (gRNA), 6 embryos, 24 cilia (AK7 KO) and 6 embryos 33 cilia (Rescue); See Figure 2B and Supp Movies 1-4). *** is < 0.0001. Graph in D is a box and whisker plot where the error bars represent the min and max values.

### Apical docking of centrioles is defective in AK7-depleted MCCs

Cilia are anchored to the apical surface of MCCs through 150-200 evenly spaced centrioles or basal bodies (Brooks and Wallingford, 2014). A decrease in the number of motile cilia could result from multiple underlying defects. To further dissect this phenotype, we performed an analysis of basal body position to determine if a defect in basal body docking was potentially driving the loss of cilia. Basal bodies are generated during a process known as centriolar biogenesis, which occurs close to the nucleus well below the apical surface. During MCC maturation, centrioles migrate apically and dock with the apical surface prior to initiating ciliogenesis (Klos Dehring et al., 2013, Zhang and Mitchell, 2015). We observed that in AK7 Crispant MCCs a considerable amount of centrioles fail to dock apically (Figure 4A, side projections). We further quantified the number of MCCs that showed sub-apical centrioles (defined as centrioles not in contact with the apical actin network). In control conditions, the occurrence of failed apical docking was negligible (4.65%in Cas9, and 7.5%in the gRNAs), indicating that these MCCs display a normal population of apically docked centrioles (Figure 4B). However, in the AK7 Crispant embryos, 46% of cells had some failed docking events, meaning that around half of the MCCs displayed some impaired centriolar maturation (Figure 4A-B). Our results suggest that failed centriole maturation and docking are contributing factors to the ciliogenesis defects, but also suggests that there are likely other contributing factors which combine to account for the significant loss of cilia in AK7 Crispants.

**Figure 4:**
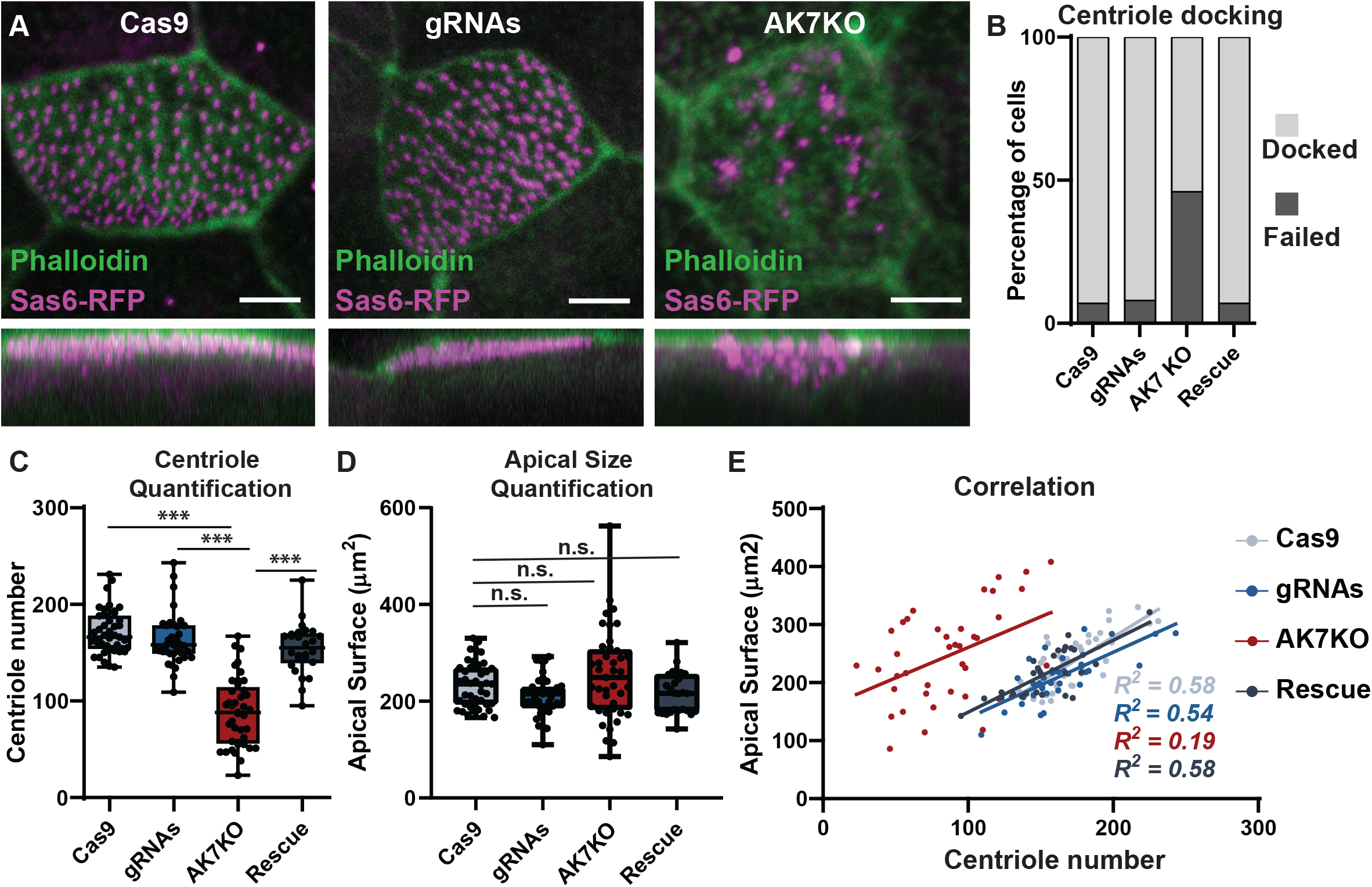
AK7 deletion causes defects in centriole biogenesis and docking. (A) Representative images of MCCs injected with Sas6-RFP (magenta) of Cas9 alone, gRNA alone or AK7 KO stained with phalloidin (green) along with side projections (lower panels). Scale is 5 μm. (B-E) Quantification of the percentage of cells with centrioles that have failed to dock (B), total centriole number (C), total apical surface area (D) and the correlation between centriole number and cell size (E) in Cas9 (n = 9 embryos, 43 cells), gRNA (n = 11 embryos, 40 cells), AK7 KO (n = 10 embryos, 41 cells) and Rescue cells (n = 8 embryos, 25 cells). *** is < 0.0001 and n.s. is not significant. Graphs in C and D are box and whisker plots where the error bars represent the min and max values.

### AK7 is an important regulator of centriole number

We next assessed the relationship between AK7 and centriole number and found that cells overexpressing (OE) AK7 had a considerable increase in centriole number. Control MCCs contained 181 +/-32 centrioles whereas in cells OE AK7 there were 234 +/-32 (p < 0.0001, Figure 5 A-B). Importantly, MCCs are known to have the ability to scale their apical area in relation to the centriole number such that they have larger apical surfaces when there are more basal bodies docked and smaller apical surfaces when less basal bodies are present (Kulkarni et al., 2021). We found that while controls cells had an average apical area of 219 +/-52 μm^2^, AK7 OE cells were significantly larger with an average size of 291+/-50 μm^2^ (p < 0.0001, Figure 5A, C). Consistent with these results when we calculated the Pearson correlation coefficient, we found similar correlations of r(35)=0.73 for control and r(35)=0.66 for AK7 OE and similar regression values (R^2^ value of 0.43 for control and 0.54 for AK7 OE, Figure 5D). Finally, we assessed overall centriole spacing by calculating the coefficient of variation of the distance between neighboring centrioles as a factor of increasing centriole number (between 2-10) and found similar variations between the two populations indicating that not only is cell size scaling with centriole number, but the organization of those centrioles is properly maintained (Figure S3).

**Figure 5:**
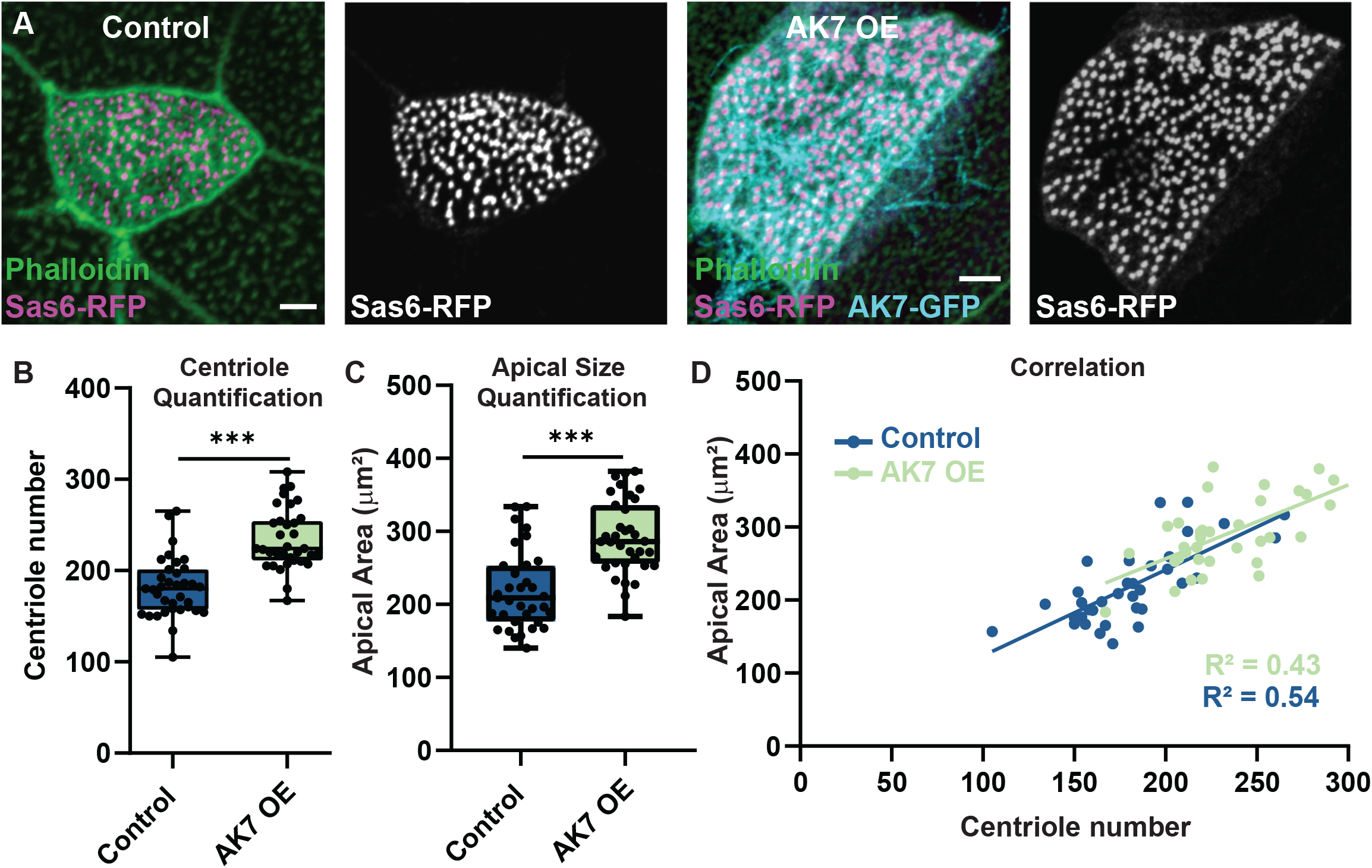
AK7 overexpression increases centriole number and cell size. (A) Representative images of embryos injected with Sas6-RFP (control, magenta and white) alone or together with AK7-GFP (AK7 OE; cyan) with phalloidin (green). Scale bar is 5 μm. (B-D) Quantification of total centriole number (B), total apical cell area (C), and the correlation between centriole number and cell size (D) in control (n = 3 embryos, 35 cells) or in AK7 OE cells (n = 4 embryos, 35 cells). *** is < 0.0001. Graphs in B and C are box and whisker plots where the error bars represent the min and max values.

Given the loss of cilia in the AK7 Crispant embryos, we next set out to characterize centriole number in AK7 Crispant MCCs. Control embryos injected with the centriolar marker Sas6-RFP, marking centrioles, together with either Cas9 alone or gRNAs alone had 171 +/- 22 and 163 +/- 26 centrioles respectively, which is consistent with previous reports for wild type embryos (Figure 4C) (Kim et al., 2021, Werner et al., 2011). In contrast, AK7 Crispants had a significant decrease in centriole number to 88 +/- 35, which could be rescued back to 154 +/- 26 with the addition of AK7-GFP mRNA (Figure 4C). Given the cell size and centriole number scaling mentioned above, we expected that AK7 Crispant MCCs would have significantly smaller apical areas given the decrease in centriole number. Surprisingly, the apical surface of AK7-depleted MCCs was not significantly different among all the conditions tested (Figure 4D). Consistent with these results we found that the correlation between apical area and centriole number was considerably lower in AK7 Cripants (r(41)=53 with an R^2^ = 0.19) compared to controls (Cas9 r(43)=0.76 with an R^2^ = 0.58, gRNA r(39)=0.74 with an R^2^ = 0.54 and rescue r(26)= 0.76 with an R^2^ = 0.58; Figure 4E). Interestingly, we also observed the spacing of centrioles to be abnormal in AK7 Crispants. The coefficient of variation was similar between wild type, Cas9, gRNA and AK7 for each number of nearest neighbors analyzed, whereas the variation was increased in AK7KO cells (Figure S3A-B). Taken together, our results suggests that AK7 has a so far uncharacterized role in regulating proper centriole number and organization, which could be a critical feature underlying the loss of mucociliary clearance.

Primary ciliary dyskinesia (PCD) is a condition characterized by impaired mucociliary clearance creating a broad range of symptoms such as respiratory dysfunction and infertility (Horani and Ferkol, 2021). To date, there are a diverse set of genes that are causative for PCD, and most of them result in immotile cilia. AK7 has been implicated in PCD (Fernandez-Gonzalez et al., 2009), but the underlying mechanistic defects have been unclear. In this study, we have identified a complex combination of phenotypes that implicate AK7 in multiple steps in proper MCC development. First, AK7 appears to be an important regulator of centriole numbers with OE leading to more centrioles and depletion leading to less centrioles. Importantly, AK7 KO cells fail to properly coordinate cell size to centriole number and have an increased variation in nearest neighbor distance suggesting that AK7 could play an important role coupling cell size to centriole biogenesis. Additionally, the centrioles that are generated in the KO embryos often fail to migrate apically and dock with the apical membrane. Interestingly, while AK7 localizes to cilia in a DPY30-dependent manner, several of its key functions occur before cilia formation (e.g. centriole number and docking) suggesting that while cilia localization may be important for some functions, AK7 has non ciliary roles as well. We propose that the combination of centriolar defects underlie the observed decrease in normal cilia number. Furthermore, when centrioles do manage to properly dock and generate a cilium, those cilia do not beat at normal frequency. Ultimately, in AK7 KO embryos the combination of a diverse range of phenotypes present as a loss of mucociliary flow that is indicative of PCD.

## Methods

All experiments in *Xenopus* embryos were performed utilizing previously described methods (Werner and Mitchell, 2013). Embryos were obtained via *in vitro* fertilization utilizing standard protocols approved by the Northwestern University Institutional Animal Care and Use Committee (Sive et al., 1998, Sive et al., 2007, Sive et al., 2010).

### Plasmids and mRNAs

AK7 of *Xenopus laevis* was amplified with PCR and cloned into pCS2+ vector containing N-terminal GFP with Enzyme/Enzyme. ∆ DPY-30 was amplified with PCR and cloned into pCS2+ containing N-terminal GFP with BamHI/Xba. SAS6 was amplified with PCR and cloned into pCS2+ containing N-terminal RFP with Enzyme/Enzyme. AK7-GFP, ∆ DPY30-GFP, and Sas6-RFP vectors were linearized with NotI and mRNA synthetized using the Sp6 mMessage Machine Kit (Life Technologies, AM1320). Embryos were injected with mRNA at 2 or 4 cells stages with 200 – 800 pg per embryo.

### CRISPR

Embryos were injected at the 1 cell stage for the CRISPR-mediated DNA editing. Embryos were injected with 500 to 750 pg of Cas9, with 500-750 pg of gRNAs (Synthego Inc), or with a combination of 500 pg of Cas9 protein and 500 pg of gRNAs for the AK7 KO condition. Single gRNAs were designed using CHOPCHOP (Labun et al., 2021). Two gRNAs were used in concert to increase the likelihood of editing. The following guides were used: AK7 Exon 1 (-strand, TTCCTGTGAATGAATCTATCTGG) and AK7 Exon 4 (+ strand, CCGAAGCAAACCTATTGATCCG).

### Immunofluorescence

Embryos were fixed at stage 28 with 4% PFA/PBS, blocked in 10% heat-inactivated goat serum, and primary and secondary antibody dilutions were prepared in 5% goat serum. Mouse anti-acetylated a-tubulin (T7451; Sigma-Aldrich) was used at 1:500 and rabbit anti-AK7 (A14600; AB Clonal) was used at 1:100. Cy-3-, Cy-5-conjugated goat anti-rabbit or goat anti-mouse, respectively, were used at 1:750 dilution. Phalloidin 405 (A30104, Invitrogen, 1:100) staining was used to visualize actin.

### Microscopy and analysis

Confocal imaging was performed on a Nikon A1R laser scanning confocal microscope using a 60X Plan-Apo objective (1.4 N.A.) Live imaging of cilia beating was performed using light microscopy on the Nikon Ti2 microscope using a 100X Plan-Apo objective (1.35 N.A.). To measure centriole number, apical area, sub-apical centrioles and cilia number, images were analyzed manually using Nikon NIS elements software. Ciliary beating frequency was analyzed with kymographs drawn on the apical-most side of cilia, over 1 second of movement (all the movies were performed at >300fps). The coefficient of variation analysis of centriole nearest neighbor distance was performed using Python script available at Github.

### Fluid flow measurement

Cilia-driven fluid flow was measured as previously described utilizing a Leica M165 FC dissecting microscope connected to the Casio Exilim 60fps mounted digital camera (Werner and Mitchell, 2013). Fluid flow velocity was calculated by determining the displacement of individual fluorescent microspheres (F8836, Invitrogen Inc.) over the surface of the embryos using a particle tracking software on Fiji (Schindelin et al., 2012).

### Statistical analysis

Normality was assayed in all data sets before any statistical analysis, through KS test, histograms and boxplot inspections. When normality was violated, non-parametric tests were used. For all histograms, bars represent the mean and error bars indicate the minimum and maximum values in the datasets. For all statistical analyses, *p < 0.05, ** p < 0.01, *** p < 0.001, and **** p < 0.0001.

## Acknowledgments

This work was supported by NIH/NIGMS to B.J.M. (R01GM089970). O.H. was supported off a diversity supplement from (R01GM119322). We want to thank Yogesh Goyal PhD and Jonas Braun for their suggestions and advice regarding the generation of analysis pipelines. We want to thank Jennifer Mitchell for critical reading and editing of the manuscript.

## Figure Legends

**Supplemental Figure 1.**
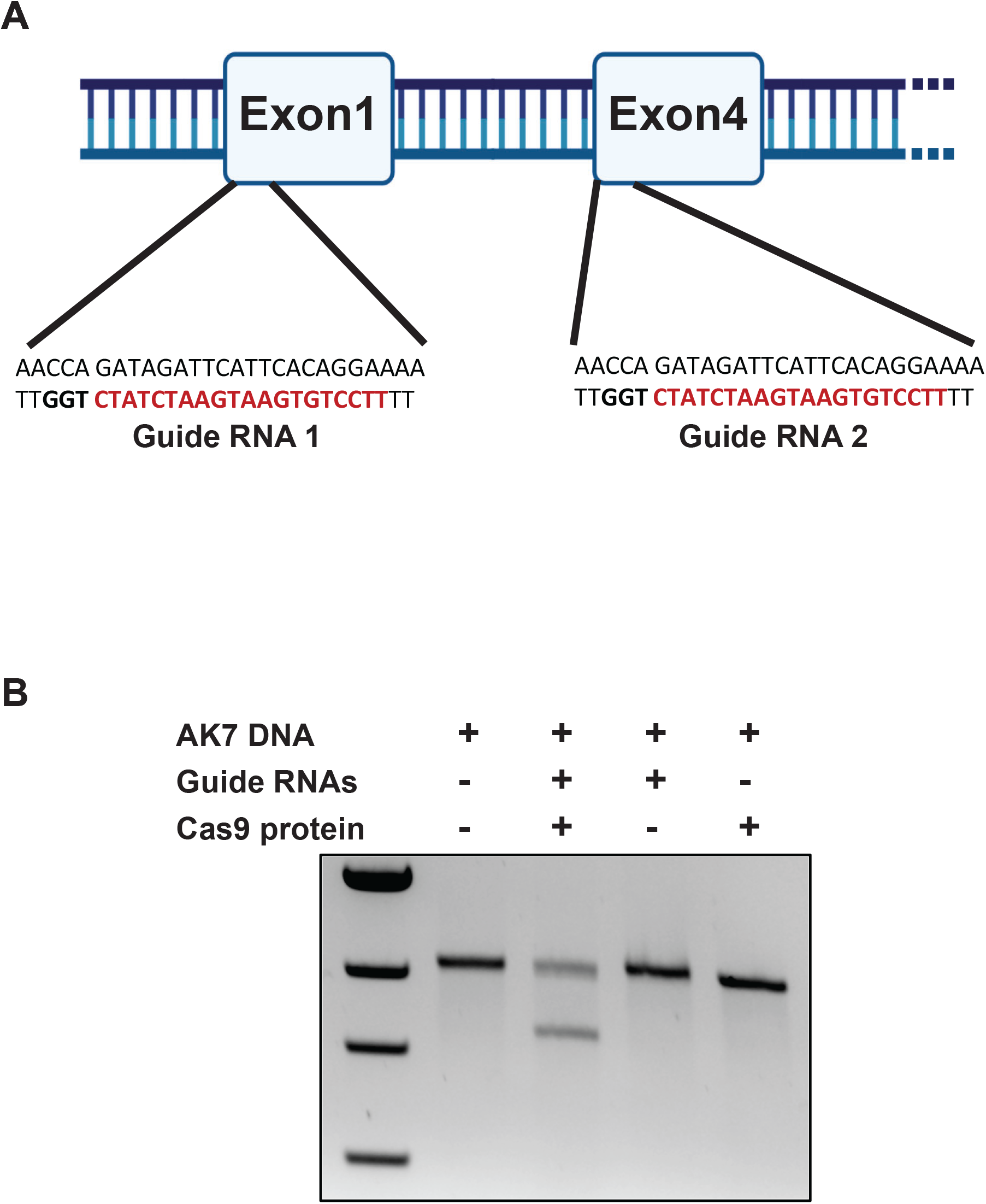
Editing capacity of AK7 via CRISPR/Cas9. (A) Schematic representation of guide RNA locations at the start of Exon1 and Exon4 together with the targeted sequences. (B) *In vitro* cleavage assay in which AK7 DNA was incubated for 1 Hr at 37º C either alone, with both guide RNAs and Cas9 protein, with the guide RNAs, or with Cas9 protein, indicating that editing only occurs in the presence of both guide RNAs and Cas9 protein.

**Supplemental Figure 2.**
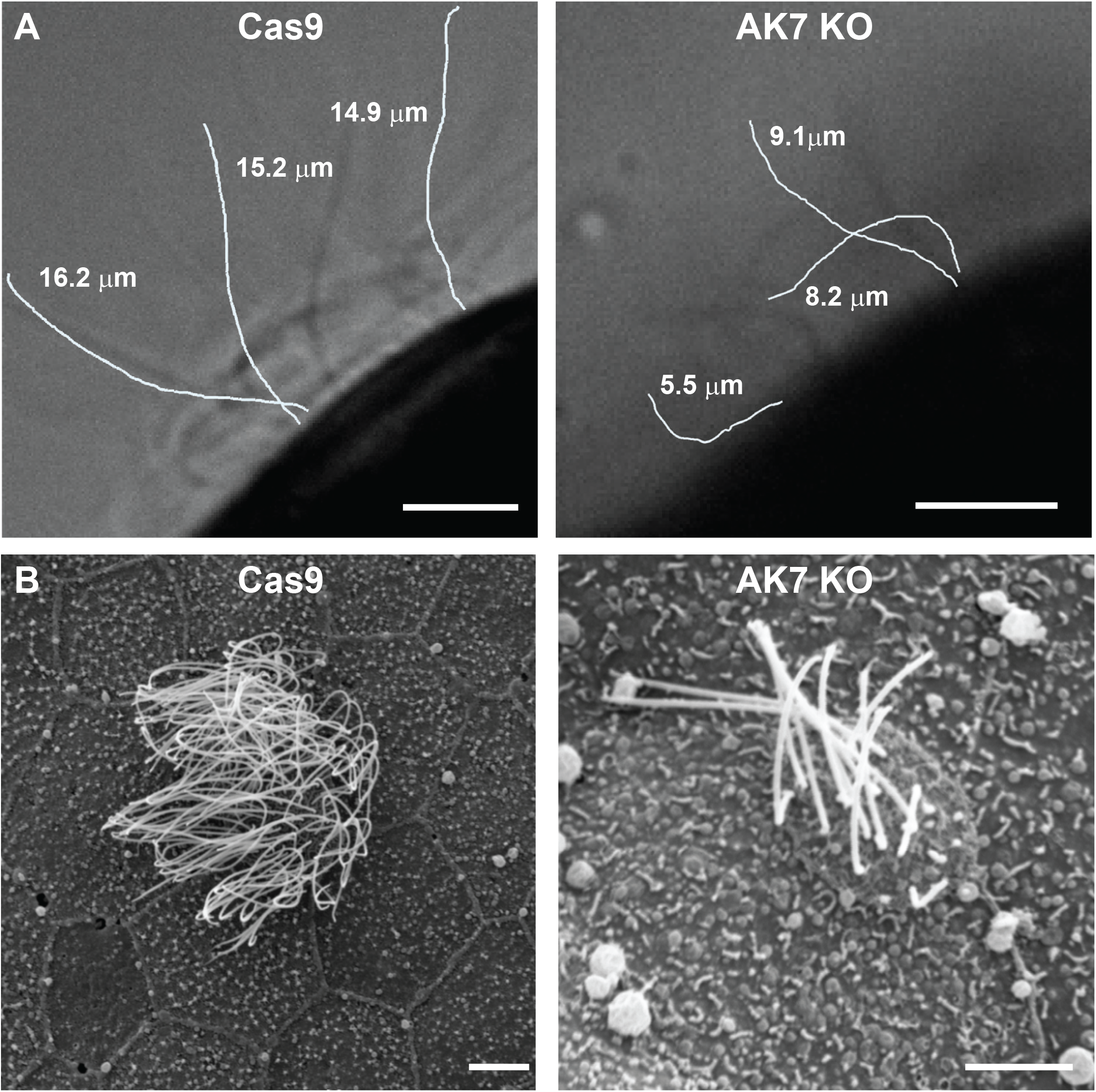
AK7 deletion affects cilia length. (A) Representative still images from high-speed movies of cilia beating (See Supplemental movies 1-4) with several cilia pseudo colored in white and labeled with their measured length. (B) Representative SEM images of Cas9 and AK7 KO cells showing the ciliogenesis defects in AK7 KO cells. Scale bar is 5 μm.

**Supplemental Figure 3.**
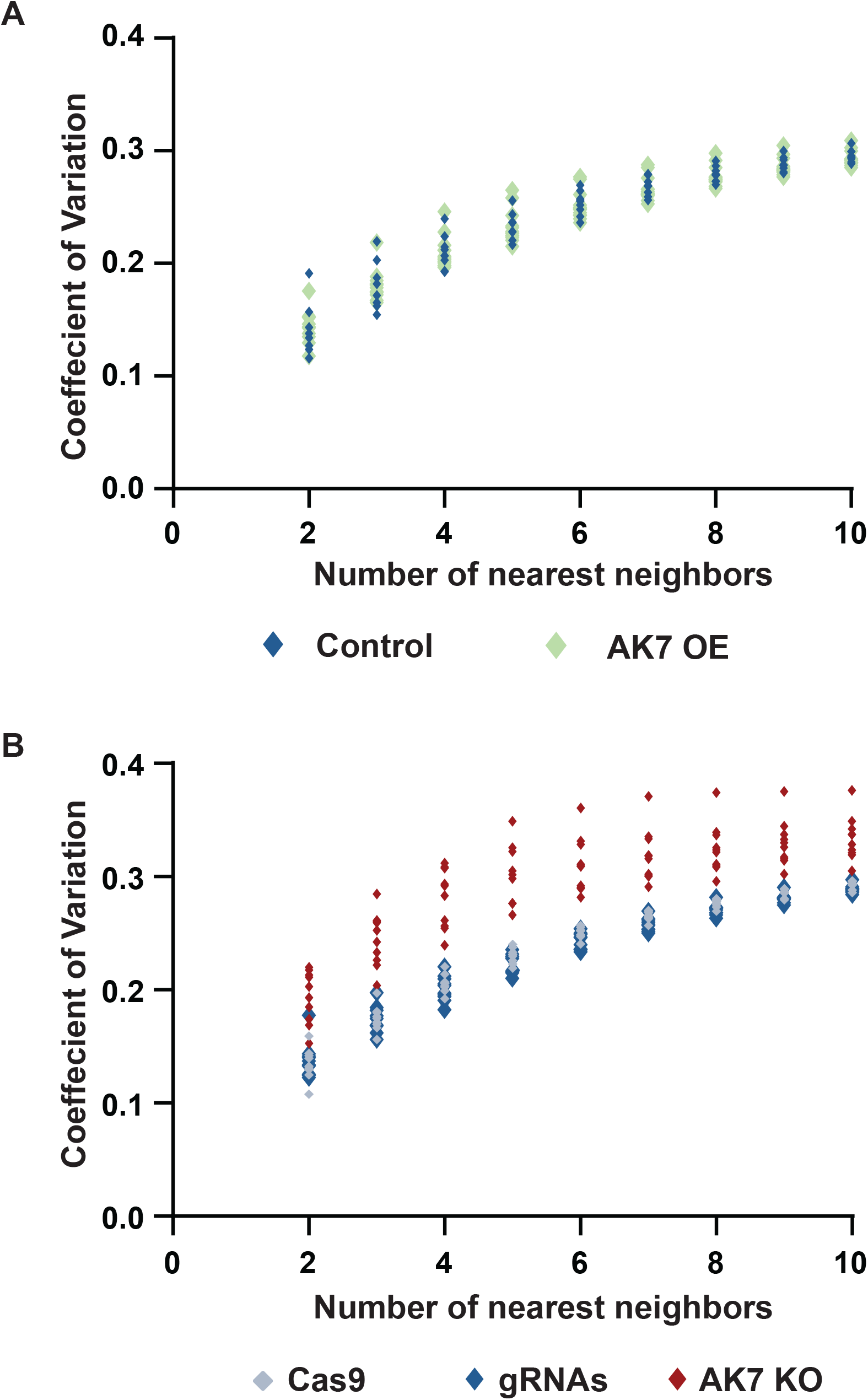
AK7 KO but not AK7 OE affects centriole spacing. (A-B) Computational analysis of the coefficient of variation for the distance between centrioles as a factor of the number of nearest neighbors analyzed. The variation is similar between control (n = 8 cells) and AK7 OE cells (n = 11 cells; (A)) whereas the variation is consistently higher in AK7 KO cells (n = 9 cells) compared to Cas9 (n = 8 cells) or gRNAs (n = 9 cells; (B)).

**Supplemental Movie 1-4. Cilia motility quantification**. Representative two second long movies of cilia beating that were used to generate kymographs for quantification (See Figure 2C-D) of Cas9 alone (Movie S1), gRNA (Movie S2), Ak7 KO (Movie S3) and Rescue cells (Movie S4).

